# ^13^C tracer analysis identifies extensive recycling of endogenous CO_2_ in vivo

**DOI:** 10.1101/2021.02.04.429777

**Authors:** Likun Duan, Daniel E. Cooper, Grace Scheidemantle, Jason W. Locasale, David G. Kirsch, Xiaojing Liu

## Abstract

^13^C tracing analysis is increasingly used to monitor cellular metabolism in vivo and in intact cells, but data interpretation is still the key element to unveil the complexity of metabolic activities. We have performed [U-^13^C]-glucose and [U-^13^C]-glutamine tracing in sarcoma-bearing mice (in vivo) and in cancer cell lines (in vitro). ^13^C enrichment of metabolites in cultured cells and tissues was determined by liquid chromatography coupled with high-resolution mass spectrometer (LC-HRMS). As expected, citrate M+2 or M+4 is the dominant mass isotopologue in vitro. However, citrate M+1 was unexpectedly the dominant isotopologue in mice receiving [U-^13^C]-glucose or [U-^13^C]-glutamine infusion. One plausible explanation is that ^13^CO_2_ produced from the oxidation of ^13^C tracers in vitro is negligible due to the dilution of HCO_3_ ^-^ supplemented to cell culture when sodium bicarbonante is used and diffusible volume of CO_2_ in the culture incubator, while endogenous ^13^CO_2_ in vivo is substantial and is fixed into the TCA cycle, purine, and serine, resulting in M+1 isotopologues. A time course study shows the generation of high abundance citrate M+1 early in plasma, which may serve as a potent non-invasive biomarker of tissue pyruvate carboxylase activity. Altogether, our results show that recycling of endogenous CO_2_ is substantial in vivo and provides important insights into the experimental design and data interpretation of ^13^C tracing assays.

## Introduction

Cellular metabolites are in dynamic homeostasis. Metabolomics is an emerging tool to measure metabolic changes that represent the dynamic status of the cell, tissue, or whole organism and facilitate a better understanding of biological processes from a global level ^1^. However, metabolomics mainly measures metabolite concentrations or the changes of metabolite levels, which reflect the net outcome of various metabolic pathways in which each metabolite is involved in, and it does not provide information on individual pathways that contribute to the production and disappearance of a specific metabolite. In contrast, stable isotope tracing provides information on metabolite dynamics and generates data on specific metabolic pathways. Stable isotope tracing has been used to probe specific metabolic pathways in various biological systems since the last century, soon after the initial isolation of isotopic tracers ^2^. Over the past decades, advances in instrumentation and data analysis software enabled fast data acquisition and determination of isotope labeling patterns ^1^. However, the data from isotope tracing experiments do not provide easily interpretable results, and interpreting ^13^C metabolite labeling patterns remains the limiting step of using stable isotope tracing to address biological questions ^3^. To extract maximal information from ^13^C tracing experiments and draw accurate conclusions, it is important to understand which labeling patterns represent different metabolic activities. Understanding the ^13^C transformations can help identify unexpected metabolic pathways of biological significance. For example, [U-^13^C]-glutamine would produce citrate M+4 following standard glutamine metabolism pathways, which involve glutaminase-mediated conversion to glutamate and then α-ketoglutarate, followed by oxidative metabolism after entering the TCA cycle ^4^. The observation of citrate M+5 suggests the existence of an alternative glutamine metabolism route, the reductive carboxylation of α-ketoglutarate, the reverse flux relative to the canonical oxidative TCA flux ^5-6^. This pathway has been demonstrated to be important for tumor growth under hypoxic conditions or tumors with mitochondrial defects and may serve as a potential therapeutic target ^5-6^. One challenge of interpreting ^13^C labeling patterns comes from the effects of culturing conditions or tissue microenvironment on cell or tissue metabolism. For instance, factors such as nutrient availability, oxygen level, or tissue context-dependent microenvironment, can greatly impact cellular metabolism ^7^ and consequently isotopic labeling of metabolites. Cells or isolated tissues fed with ^13^C tracers ex vivo also tend to have different labeling patterns from the same type of tissues of animals receiving in vivo tracing ^1, 8^. These factors result in difficulties in data interpretation and developing new approaches to decipher labeling patterns.

Pyruvate carboxylase, a mitochondrial enzyme that catalyzes the production of oxaloacetate by combining pyruvate and bicarbonate, is one of the major anaplerotic enzymes. Pyruvate carboxylase supports gluconeogenesis in the liver and kidney by providing oxaloacetate. Pyruvate carboxylase is important for de novo synthesis of fatty acid in adipocyte tissues and is also involved in de novo synthesis of a neurotransmitter, glutamate, in astrocytes. Studies have shown that pyruvate carboxylation is required for tumor growth when glutamine metabolism is suppressed ^9-10^. ^13^C tracing has been used to study TCA anaplerosis in vitro and in vivo and citrate M+3 has been used to monitor pyruvate carboxylase activity after [U-^13^C]-glucose tracing ^10-11^. For instance, under cell culture conditions, pyruvate M+3 can enter the TCA cycle via pyruvate carboxylase and lead to the appearance of citrate M+3. Indeed, citrate M+3 is observed in cultured cells fed with [U-^13^C]-glucose. Nevertheless, citrate M+1 has been reported to be more abundant than M+3 in lung cancer, liver, and kidney after continuous infusion of [U-^13^C]-glucose or dietary delivery of [U-^13^C]-glucose in vivo ^8, 11^, but it remains poorly understood why M+1 is more abundant in vivo.

Here we report the distinct ^13^C labeling patterns of TCA intermediates in cultured cells, in-vivo tumors, and non-tumor tissues after [U-^13^C]-glucose or [U-^13^C]-glutamine tracing. We identified M+1-labeling of TCA intermediates are the most abundant species in vivo, but not in vitro. This finding is consistent with the results of in vivo [U-^13^C]-glucose tracing performed by other investigators ^8, 11-13^. We hypothesized that endogenous CO_2_ is labeled by ^13^C tracers and subsequently used to produce M+1 isotopologues of TCA metabolites. To test this hypothesis, we traced the product of other CO_2_-consuming reactions such as purine biosynthesis and measured adenosine M+1 in tumor and non-tumor tissues. We then compared the time course of M+1 and M+3 TCA intermediates and observed that M+1 citrate appeared earlier than M+3, which further suggests that CO_2_ is produced endogenously and fixed into the TCA cycle. These findings provide a new paradigm to understand carbon atom transformations in vivo and should be taken into account when developing mathematic models to better reflect carbon flux.

## Results

### M+1-labeling of TCA metabolites is dominant in the in vivo tumor but not in in vitro cultured cells fed with ^13^C glutamine tracer

To investigate the ^13^C labeling patterns of TCA intermediates in cancer cells (in vitro), and in animals (in vivo) with [U-^13^C]-glutamine, two different types of tracing experiments were performed (Fig.1A). Sarcoma cells, generated from primary mouse sarcomas, were incubated in RPMI 1640 containing 2 mM [U-^13^C]-glutamine and 10% dialyzed FBS. Intracellular metabolites were extracted after 3 hours of tracing and subsequently analyzed using LC coupled with a high resolution mass spectrometer (LC-HRMS). Sarcoma-bearing mice were infused with [U-^13^C]-glutamine for 3 hours. At the end of infusions, sarcoma samples were collected and analyzed using LC-HRMS. ^13^C enrichment was calculated based on metabolite peak area. Natural abundance correction was performed using software R with Bioconductor R package IsocorrectionR (21). In tumor (in vivo), [U-^13^C]-glutamine was converted into [^13^C_5_]-2-oxoglutarate (M+5) via glutaminolysis (Fig. 1B), followed by oxidative decarboxylation of 2-oxoglutarate M+5, resulting in succinate-CoA and succinate M+4. The M+4 carbon chain remains labeled in the next two reactions to produce malate M+4. Malate M+4 will be oxidized to oxaloacetate M+4, which is then used to generate citrate M+4. The citrate M+4 was then dehydrated by aconitase to give M+4 cis-aconitate. Isocitrate dehydrogenase (IDH) catalizes the reversible reaction of isocitrate to alpha-ketoglutarate and CO_2_, and the reverse reaction contributes to the generation of citrate M+5 and cis-aconitate M+5 (Fig. 1B). In the cultured sarcoma cells, M+4 and M+5 isotopologues are the major species as expected. Surprisingly, unlike the labeling patterns in cultured cells, M+1-labeled intermediates are the major isotopologues in mouse sarcomas in vivo. A plausible explanation is that decarboxylation of [U-^13^C]-glutamine produces ^13^CO_2_, which is subsequently fixed into the TCA cycle in vivo. This model challenges the common assumption that endogenous ^13^CO_2_ from decarboxylation of [U-^13^C]-glutamine is diluted by the body pool of bicarbonate and thus the contribution of ^13^CO_2_ to ^13^C-labeling is not significant. Furthermore, TCA reactions are often referred to as having a net flux in the clockwise direction. The observation of M+1-labeled succinate, fumarate, and malate indicates the reverse reactions and exchange fluxes from oxaloacetate to succinate.

**Figure 1:**
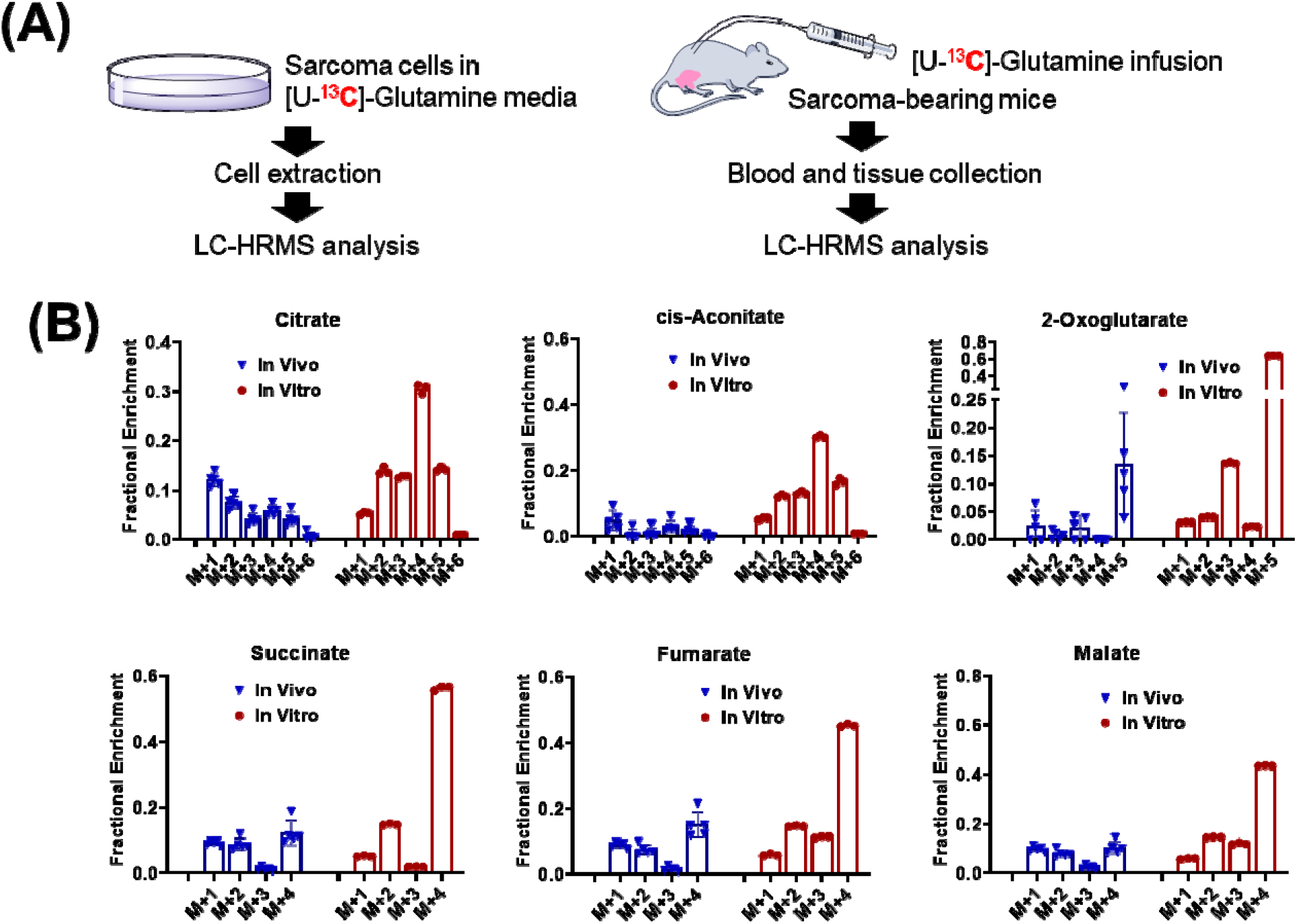
Distinct ^13^C labeling of TCA intermediates with [U-^13^C]-glutamine is observed in in vitro cell cultures and in tumors in vivo. (A) Schematic of in vivo and in vitro tracing anaylsis. (B) ^13^C labeling patterns of metabolites from sarcoma cells (in vitro) fed with [U-^13^C]-glutamine medium for 3 hours (hrs) and sarcoma tissue from sarcoma-bearing mice (in vivo) receiving [U-^13^C]-glutamine infusion for 3 hrs. Data are presented as mean ± S.D. n=5 for animals and n=3 for cultured cells.

### M+1-labeled TCA metabolites by [U-^13^C]-glucose is dominant in the in-vivo tumor but not in cultured cells

To investigate whether glucose-drived CO_2_ can also be recycled, we next performed in vitro and in vivo tracing experiments using [U-^13^C]-glucose (Fig. 2A). Through glycolysis, [U-^13^C]-glucose is converted into [^13^C_3_]-pyruvate (M+3), which then feeds a series of reactions in the TCA cycle. Pyruvate M+3 is first converted into acetyl-CoA, and the M+2 acetyl group of acetyl-CoA is then transferred to the four-carbon oxaloacetate to form citrate M+2 (Fig. 2B). The citrate M+2 is then dehydrated by aconitase to give cis-aconitate M+2, which subsequently goes through a series of chemical transformations to yield α-ketoglutarate M+2, losing one carboxyl group as CO_2_. After oxidative decarboxylation of α-ketoglutarate M+2, a four-carbon chain is generated as succinyl-CoA, which is further converted into succinate M+2. The second carbon lost as CO_2_ is originated from the carbon of oxaloacetate. The succinate M+2 remains labeled in the next two enzymatic reactions to provide malate M+2. During the second round of the TCA cycle, oxaloacetate M+2 reacts with acetyl-CoA M+2 and produces citrate M+4. Indeed, M+4 was detected in cutured sarcoma cells (in vitro) and in sarcomas in mice (in vivo) and M+4 is less abundant than M+2. Surprisingly, M+1 species are the dominant isotopologues for all the TCA metabolites in the tumor but not in the cultured cancer cells. Taken together, the appearance of high abundant M+1 TCA intermediates in [U-^13^C]-glutamine (Fig. 1) and [U-^13^C]-glucose tracing (Fig. 2) experiments can be explained by the recycling of endogenous ^13^CO_2_.

**Figure 2:**
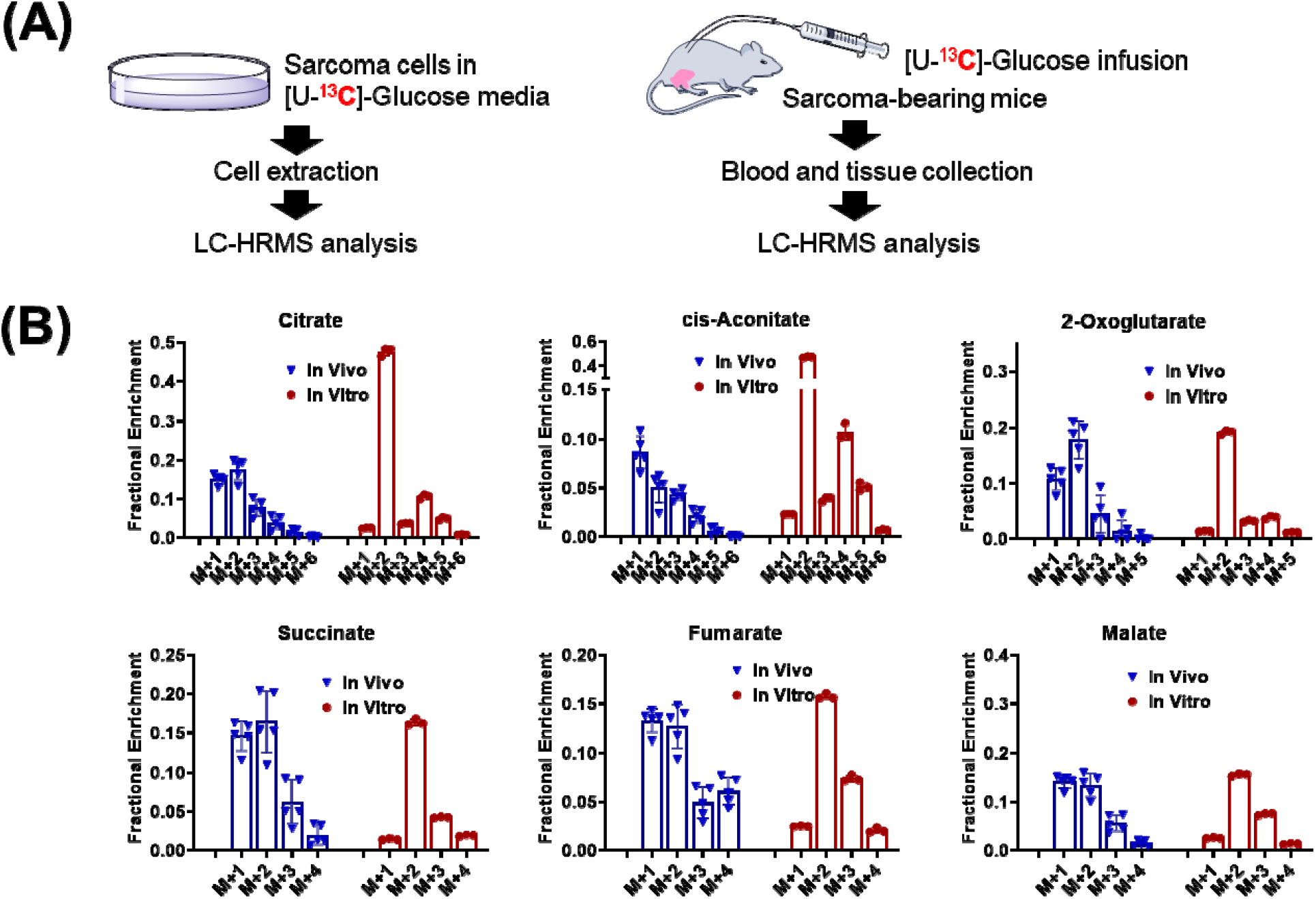
Distinct ^13^C labeling of TCA intermediates with [U-^13^C]-glucose is observed in in vitro cell cultures and in tumors in vivo. (A) Schematic of in vivo and in vitro tracing assays. (B) ^13^C labeling patterns of metabolites from sarcoma cells (in vitro) fed with [U-^13^C]-glucose medium for 3 hours (hrs) and sarcoma tissue from sarcoma-bearing mice (in vivo) receiving [U-^13^C]-glucose infusion for 3 hrs. Data are presented as mean ± S.D. n=5 for animals and n=3 for cultured cells.

### M+1-labeled TCA intermediates is also observed in non-tumor tissues in vivo

To investigate whether CO_2_ recyling is a tumor-specific metabolic event or not, we further looked into the labeling patterns of TCA intermediates in non-tumor tissues.

Similar to what we observed in tumor, citrate M+1, succinate M+1, and malate M+1 are dominant isotopologues in mouse liver and skeletal muscle (Fig. 3A). In fact, a similar result of TCA intermediate enrichment in normal tissues after infusing mice with [U-^13^C]-glucose was reported by other independent investigators^13^, but the source of the M+1 species was not identified. We downloaded the relevant source data from the supplementary material of this paper and analyzed the labeling pattern of citrate, succinate, and malate in different normal tissues (Fig. 3B). For citrate, the encrichment fractions of M+1 and M+2 are comparable. For succinate and malate, M+1 is even more abundant than M+2. These results indicate that endogenously generated ^13^CO_2_ participates in anaplerotic metabolism in non-tumor tissues as well.

**Figure 3:**
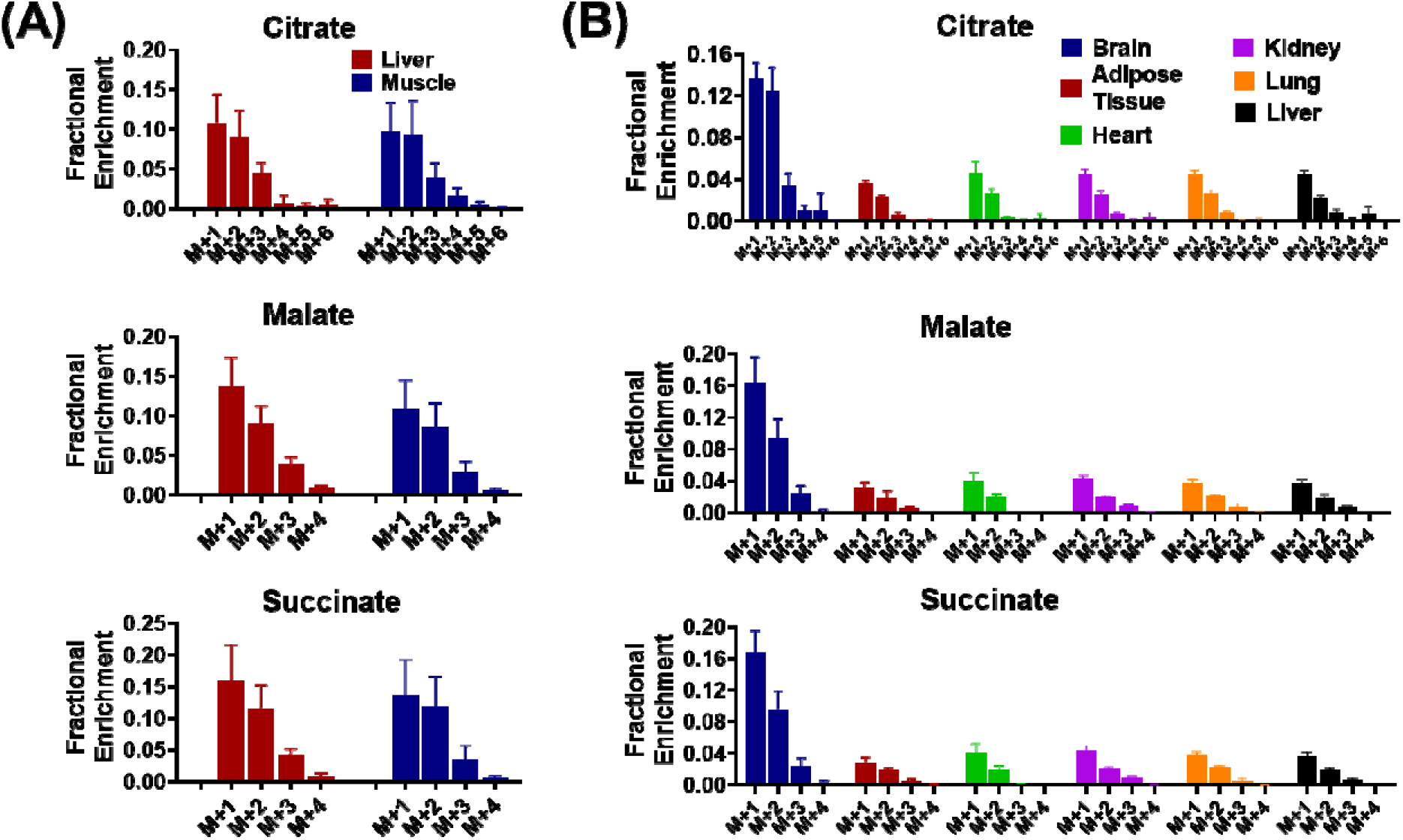
M+1-labeled TCA intermediates are observed in non-tumor tissues in vivo. A) ^13^C labeling patterns of metabolites in skeletal muscle and liver from sarcoma-bearing mice receiving [U-^13^C]-glucose infusion for 3 hours. (B) ^13^C labeling patterns of metabolites in different types of organs (data from Hui et al., Nature 2017)^13^. Data are presented as mean ± S.D. n=5 animals per group.

### Endogenous CO_2_ contributes to purine biosynthesis in vivo

To explore whether CO_2_ recycling participates in other metabolic events, we then looked into de novo purine synthesis, which consumes CO_2_ (Fig. 4A). An intermediate in purine biosynthesis, aminoimidazole ribonucleotide (AIR) is combined with ^13^CO_2_ by AIR carboxylase to produce CAIR M+1. The CAIR was then converted into (S)-2-[5-Amino-1-(5-phospho-D-ribosyl)imidazole-4-carboxamido]succinate (SAICAR) by SAICAR synthetase, followed by several concerted steps to yield inosinic acid M+1. The inosinic acid M+1 is then converted by 5’-ribonucleotide phosphohydrolase into inosine M+1, which is subsequently converted into adenosine M+1 by adenosine aminohydrolase.^13^C labeling patterns of adenosine from sarcoma cells fed with [U-^13^C]-glutamine/[U-^13^C]-glucose in vitro and from sarcomas of sarcoma-bearing mice receiving [U-^13^C]-glutamine/[U-^13^C]-glucose infusion in vivo for 3 hours are shown in Fig. 4B. No substantial labeling of adenosine with [U-^13^C]-glutamine was observed in cultured cells. However, adenosine M+1 was detected in sarcomas from sarcoma-bearing mice receiving the [U-^13^C]-glutamine infusion, strongly suggesting that ^13^CO_2_ produced from [U-^13^C]-glutamine participates in purine biosynthesis. The [U-^13^C]-glucose tracing leads to M+5 labeled-adenosine in both cultured cells and tumor, indicating the successful labeling of ribose 5-phosphate (R5P) by [U-^13^C]-glucose through the pentose phosphate pathway in vitro and in vivo. However, M+1-labeling of adenosine with [U-^13^C]-glucose was only observed in the tumor in vivo. The observation of adenosine M+1 in sarcomas after [U-^13^C]-glutamine or [U-^13^C]-glucose infusion in mice but not in cultured sarcoma cells is consistent with our previous observations that ^13^CO_2_ recycling is substantial in vivo but not in vitro (Figs. 1-2).

**Figure 4:**
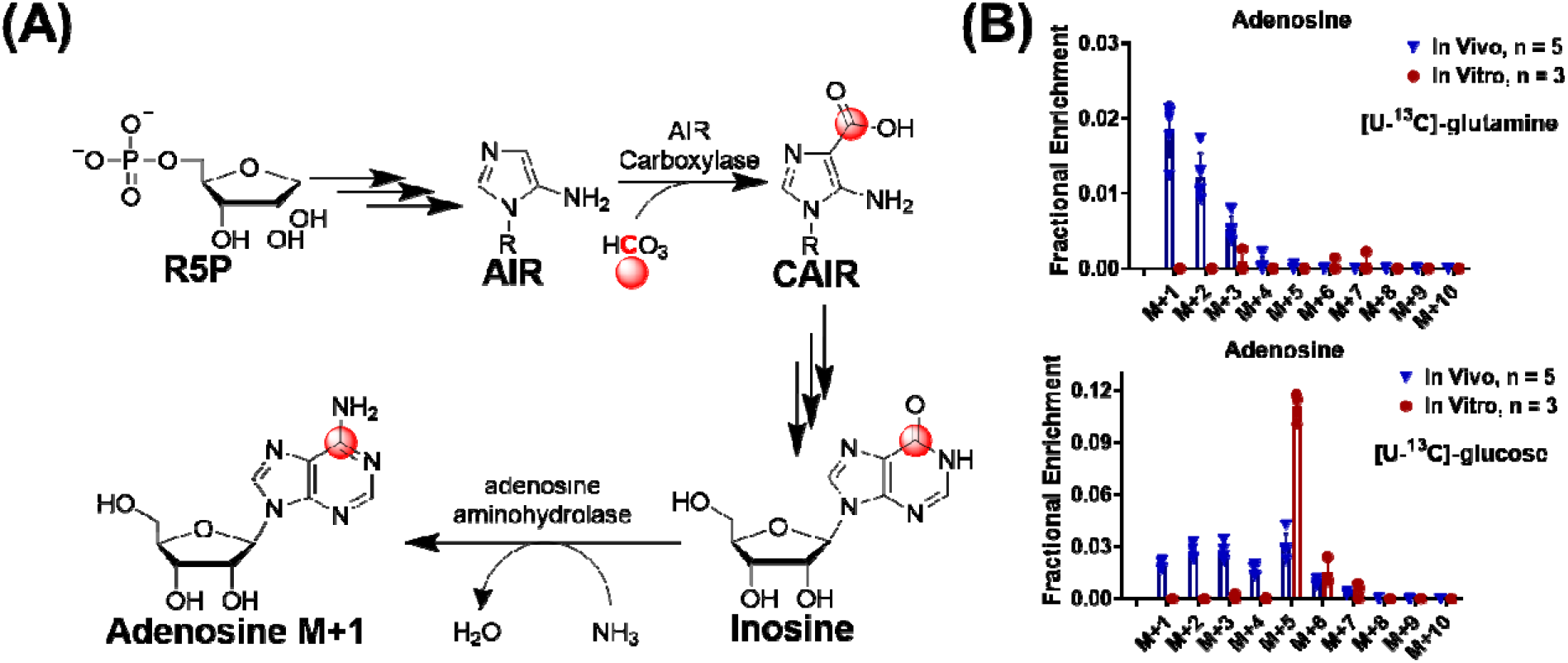
^13^CO_2_ contributes to adenosine biosynthesis. (A) Schematic of adenosine biosynthesis from various carbon sources. (B) ^13^C labeling pattern of adenosine from sarcoma cells fed with [U-^13^C]-glutamine/[U-^13^C]-glucose in vitro for 3hours and labeling patterns of adenosine in sarcomas from sarcoma-bearing mice receiving [U-^13^C]-glutamine/[U-^13^C]-glucose infusion in vivo for 3 hours. Data are presented as mean ± S.D. n=5 for animals and n=3 for cultured cells. (R5P, Ribose 5-phosphate; AIR, Aminoimidazole ribotide; CAIR, 5-Amino-1-(5-phospho-D-ribosyl)imidazole-4-carboxylate.

### Endogenous CO_2_ contributes to serine biosynthesis in vivo

We further investigated other ^13^C labeling patterns which can be explained by ^13^CO_2_ recycling. In the [U-^13^C]-glutamine tracing experiment, serine M+1 was observed in mouse sarcomas in vivo but not in sarcoma cells cultured in a CO_2_ incubator (Fig. 5). Oxaloacetate M+1 is generated by incorporating the ^13^CO_2_ with nonlabelled pyruvate (Fig. 5A). Oxaloacetate M+1 is then converted into PEP M+1 by introducing a phosphate group from ATP, catalyzed by PEPCK-C. PEP M+1 is then hydrolyzed by enolase to produce 2PG M+1, which is in equilibrium with 3PG M+1 in the presence of PGAM. 3PG M+1 is eventually converted into serine M+1. Indeed, consistent with the model that ^13^CO_2_ recycling is substantial in vivo, 2PG/3PG M+1 was only observed in tumors after in vivo infusion, but not in cultured tumor cells (Fig. 5B). Subseqently, de novo serine biosynthesis from 3PG M+1 leads to the production of serine M+1 (Fig. 5B). These results suggest that ^13^CO_2_ participates in the biosynthesis of serine in vivo.

**Figure 5:**
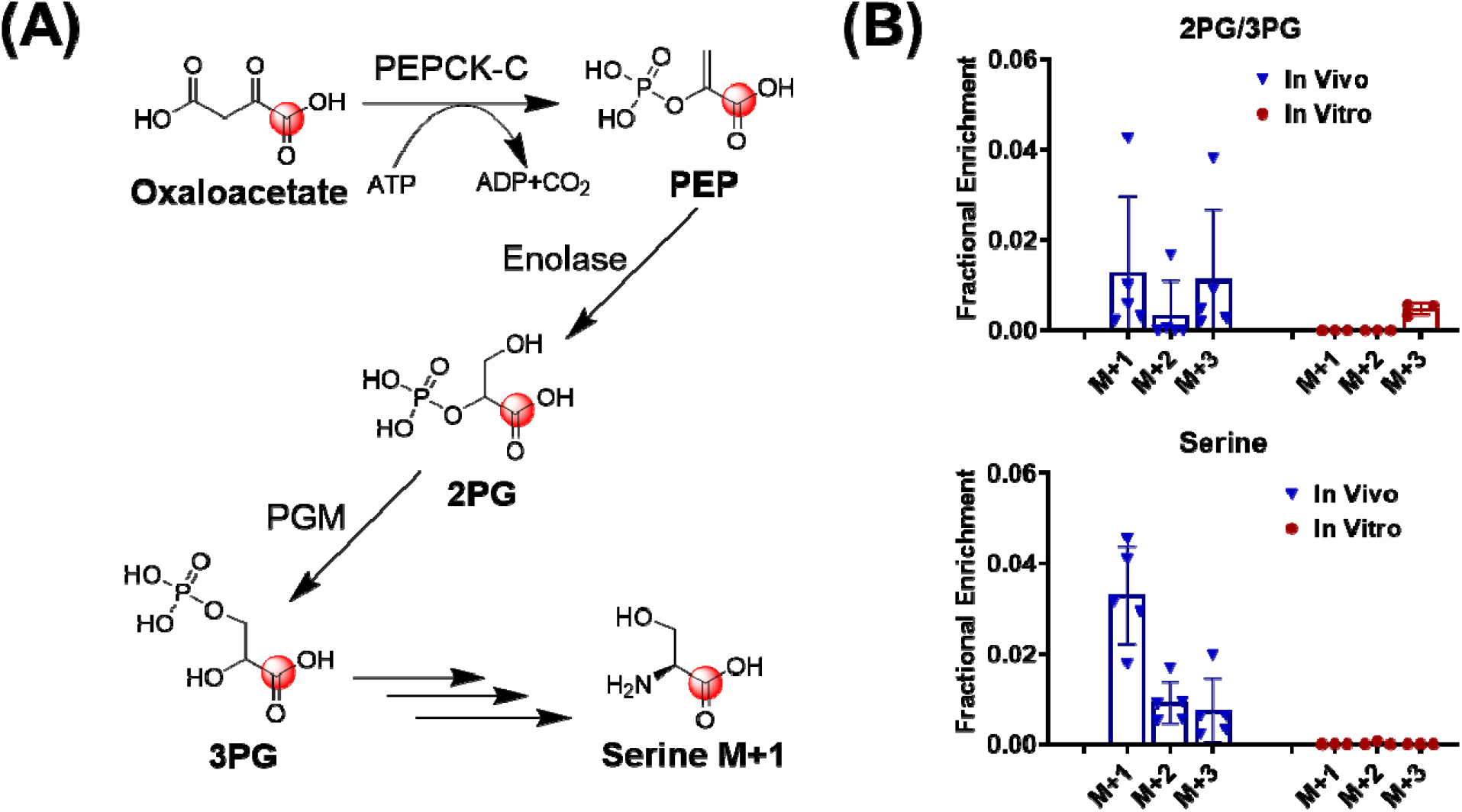
^13^CO_2_ contributes to serine biosynthesis. (A) Schematic of ^13^C incorporation into serine via the serine biosynthesis pathway. (B) ^13^C labeling pattern of 2PG/3PG and serine from sarcoma cells fed with [U-^13^C]-glutamine in vitro for 3hours and labeling patterns of 2PG/3PG and serine in sarcomas from sarcoma-bearing mice receiving [U-^13^C]-glutamine infusion in vivo for 3 hours. Data are presented as mean ± S.D. n=5 for animals and n=3 for cultured cells. (PEP, Phosphoenolpyruvate; PEPCK-C, phosphoenolpyruvate carboxykinase; 2PG, 2-Phosphoglyceric acid; 3PG, 3-Phosphoglyceric acid; PGM, 2,3-bisphosphoglycerate-dependent phosphoglycerate mutase).

### Citrate M+1 is a potent marker of pyruvate carboxylase in vivo

Pyruvate anaplerosis is the counterpart to glutamine anaplerosis and allows the TCA cycle to continuously oxidize acetyl-CoA simultaneously to provide carbon backbones for biomass production. In the [U-^13^C]-glucose tracing experiment, [^13^C_3_]-pyruvate (M+3) is produced through glycolysis. Pyruvate M+3 enters the TCA cycle through pyruvate dehydrogenase or pyruvate carboxylase, leading to the production of M+2 acetyl-CoA or M+3 oxaloacetate, respectively, which subsequently contributes to the production of citrate M+2 and M+3, respectively. Hence, citrate M+3 is usually used as a marker of pyruvate carboxylase in many studies ^1^. Surprisingly, the time-course experiment (Fig. 6) demonstrated that citrate M+1 is more abundant than M+3 and also appears earlier than M+3 in the liver, skeletal muscle and plasma of mice receiving [U-^13^C]-glucose tracing. Non-labeled pyruvate and [U-^13^C]-glucose-derived ^13^CO_2_ produce oxaloacetate M+1 via pyruvate carboxylase. Oxaloacetate M+1 is then converted into citrate M+1. Citrate M+1 mirrors that in tissues (liver and skeletal muscle), and thus plasma citrate M+1 may serve as a useful biomarker of pyruvate carboxylase.

**Figure 6:**
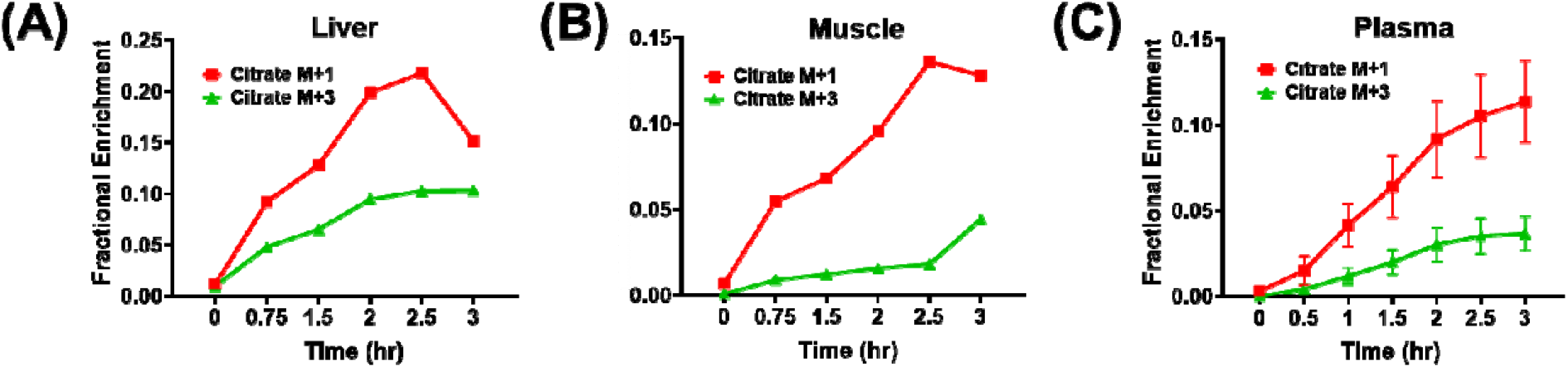
Plasma citrate M+1 is a potent biomarker of pyruvate carboxylase activity. Time course of citrate M+1 and M+3 in liver (A), skeletal muscle (B), and plasma (C) of mice receiving [U-^13^C]-glucose infusion for 3 hours (hrs). Data in (C) are presented as mean ± S.D. n=5 animals per group for the plasma sample only.

When taken together, these results suggest that even though ^13^CO_2_ produced from the decarboxylation of ^13^C tracers is negligible in vitro, endogenous ^13^CO_2_ produced in vivo is substantial and subsequently fixed into the TCA cycle, purine, and serine, resulting in M+1 isotopologues in different metabolic pathways, which are summarized in Fig. 7.

**Figure 7:**
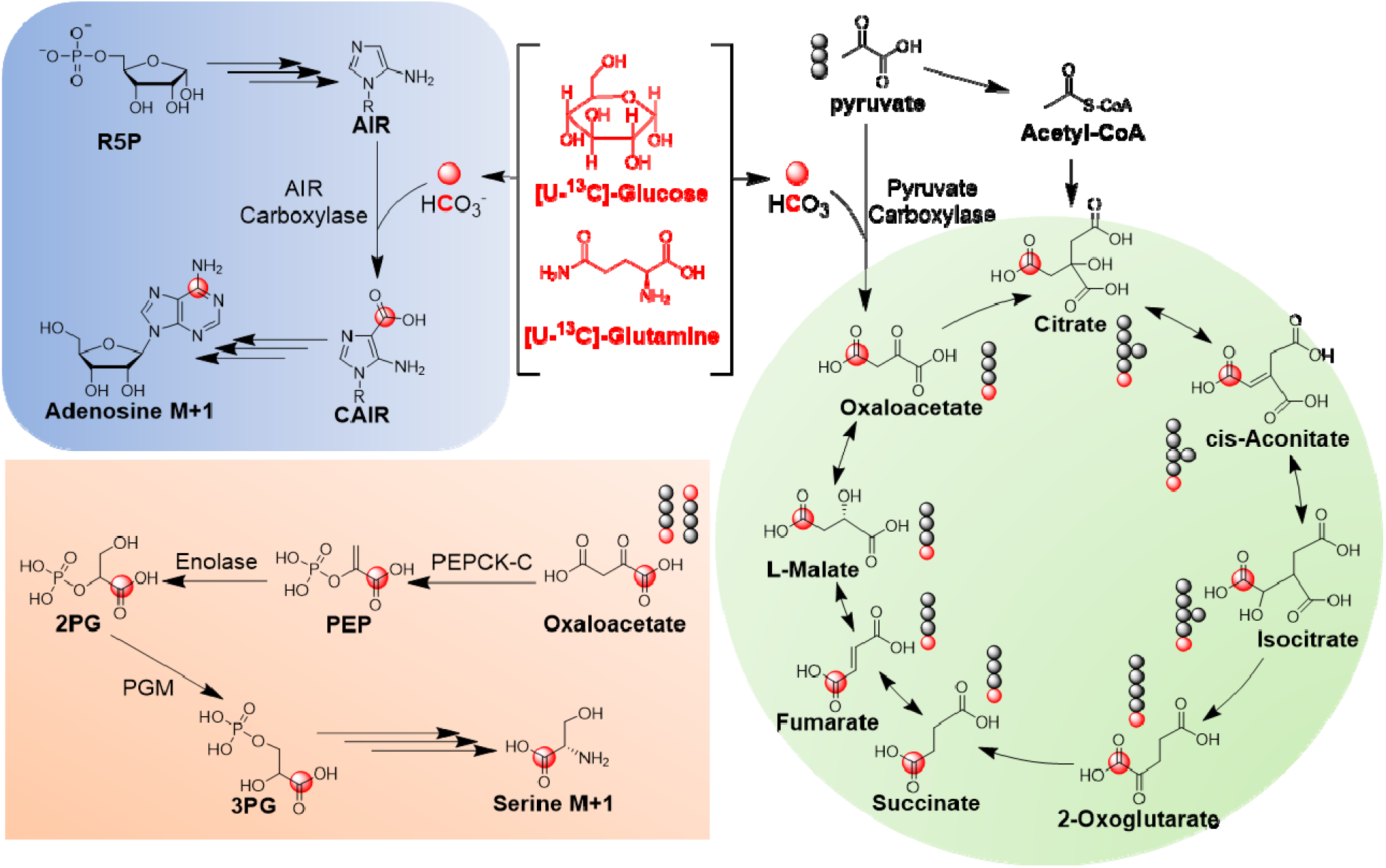
Proposed route of CO_2_ incorporation into different metabolomic pathways.

## Discussion

Compared to metabolomics which detects changes in metabolite levels, stable isotope tracing is more informative, because it provides activity readout of metabolic enzymes in intact cells or whole organisms. Precisely relating metabolite enrichment patterns to a specific metabolic enzyme or metabolic pathway requires in-depth knowledge of metabolic reactions and biological systems in which the isotope tracing assay is performed. We performed [U-^13^C]-glutamine and [U-^13^C]-glucose tracing in sarcoma cells cultured in standard in vitrocell culture conditions and also in vivo in mice. We observed distinct enrichment patterns of TCA intermediates, featured by high-abundant M+1 species in mice, but not in cultured cells. Interestingly, observing high-abundant M+1 species in vivo recapitualtes previously published results by other groups ^8, 11, 13-14^, but the reason for the difference in the M+1 species between in vivo and in vitro experiments has not previously been explained. Our results are consistent with a model where the difference in M+1 species is due to the different sources of CO_2_ in vivo and in vitro. Endogenous CO_2_ in cell culture is diluted by exogenous CO_2_ and NaHCO_3_ from the artificial conditions within an incubator, and hence the ^13^C enrichment by [U-^13^C]-glutamine or [U-^13^C]-glucose is negligible. In contrast, CO_2_ from endogenous metabolism is substantial in mice and our results indicate that CO_2_ is labeled in the presence of [U-^13^C]-glucose or [U-^13^C]-glutamine. ^13^CO_2_ re-enters the TCA cycle via pyruvate carboxylase-mediated pyruvate carboxylation, resulting in M+1-labeled TCA intermediates. High-abundant M+1 TCA species in multiple tissues of mice receiving [U-^13^C]-glucose were also observed by other groups. [U-^13^C]-glutamine may contribute to ^13^CO_2_ production via α-ketoglutarate dehydrogenase- and isocitrate dehydrogenase-mediated decarboxylation reactions. [U-^13^C]-glucose may contribute to ^13^CO_2_ production via oxidative pentose phosphate pathway or pyruvate dehydrogenase-mediated pyruvate decarboxylation. These findings suggest that special attention needs to be paid when M+1 species are used to track specific pathways. For example, [1-^13^C]-gltuamine can be used as a tracer in cancer cells where citrate M+1 is used to monitor reductive glutaminolysis activity. This approach is acceptable when it is used in vitro because the endogenous CO_2_ contribution is negligible in the presence of an exogenous CO_2_ supplement. However, our findings reveal that this approach should be used with caution to estimate reductive glutaminolysis in vivo, because [1-^13^C]-glutamine will label CO_2_, and citrate M+1 may come from pyruvate carboxylation and/or reductive glutaminolysis. In addition, adenosine M+1 may be misinterpreted as the contribution via one-carbon metabolism in vivo, but [U-^13^C]-glutamine is not likely contributing to one-carbon metabolism, so adenosine M+1 is also due to recycling ^13^CO_2_.

Replenishing the TCA cycle (a process termed anaplerosis) is important to maintain homeostasis. The carbon source of anaplerosis varies from tissue to tissue and is also dependent on physiological conditions (e.g., fast vs fed state). Pyruvate carboxylase has been identified to play an important role in regulating anaplerosis ^11^. Hence, pyruvate carboxylase is involved in various types of diseases and is an emerging therapeutic target. Pyruvate carboxylase deficiency is a genetic disease present at birth, leading to damage to tissues and organs. Pyruvate carboxylase is also involved in tumorigenesis ^10, 15-17^. High pyruvate carboxylase was shown to be correlated with glycemia ^18^. Tissue-specific inhibition of pyruvate carboxylase reduced plasma glucose concentrations and could be a potential therapeutic approach for nonalcoholic fatty liver disease, hepatic insulin resistance, and type 2 diabetes ^18^. Citrate M+3 is often used to reflect pyruvate carboxylase activity in [U-^13^C]-glucose tracing experiments, but our results show that citrate M+1 is more abundant than citrate M+3 in both tissues and plasma. Hence, citrate M+1 in plasma may be a better readout of tissue pyruvate carboxylase and may be useful to monitor the efficacy of treatments targeting pyruvate carboxylase.

Furthermore, CO_2_ recycling may also affect the interpretation of the respiratory quotient (RQ), which is the ratio of CO_2_ production to O_2_ consumption. For every mole of glucose undergoing complete oxidation, 6 moles of CO_2_ are produced and 6 moles of O_2_ are consumed, so the RQ is 1. Each mole of palmitate produces 16 moles of CO_2_ and consumes 23 moles of O_2_, so the RQ is close to 0.7 when palmitate is completely oxidized. Hence, the RQ is used as an indicator of energy fuel (e.g. carbohydrate or fat) being metabolized in the body and helps to plan nutritional therapy. The RQ is determined by comparing exhaled gases to room air, but this approach does not take in vivo CO_2_ recycling into consideration. Therefore, the RQ will be underestimated when CO_2_ recycling is substantial, resulting in an inaccurate interpretation of fuel source utilization.

## Conclusion

In summary, understanding enrichment patterns and individual isotopologue provides rich information on metabolic reactions. The production and recycling of ^13^CO_2_ from the decarboxylation of [U-^13^C]-glucose or [U-^13^C]-glutamine is negligible in vitro due to dilution by the exogenous HCO_3_ ^-^/CO_2_ source, but in vivo the incorporation of endogenous ^13^CO_2_ into M+1 TCA intermediates and other metabolites is substantial and should be considered. M+1 TCA intermediates in vivo also suggest the isotopic exchange of oxaloacetate, malate, fumarate, and succinate, indicating the reversibility of malate dehydrogenase, fumarase, and succinate dehydrogenase. These findings not only provide an interpretation of distinct labeling patterns of TCA intermediates in vivo and in vitro but also provide insights into the proper design of ^13^C tracing experiments and modeling when ^13^C tracers are used to study in vivo metabolism. In addition, using M+1 citrate in plasma as a readout of tissue pyruvate carboxylase potentially offers a noninvasive approach for the diagnosis of diseases of pyruvate carboxylase deficiency or monitoring of treatment efficacy when targeting pyruvate carboxylase is employed as a therapeutic strategy for the treatment of diseases.

## Experimental procedures

### Reagents

Optima LC-MS grade of ammonium acetate, water, acetonitrile and methanol were purchased from Fisher Scientific. [U-^13^C_6_]-glucose and [U-^13^C_6_]-glutamine were obtained from Cambridge Isotope Laboratories. RPMI 1640 medium and Fetal Bovine Serum (FBS) were obtained from Thermo Fisher Scientific. Dialyzed FBS was obtained from Thermo Fisher Scientific. Jugular vein catheters, vascular access buttons, and infusion equipment were purchased from Instech Laboratories.

### Cell culture

Mouse primary sarcoma cell lines were generated from Pax7CreER-T2, p53^FL/FL^, LSL-Nras^G12D^ tumors as previously described ^19^. Mouse sarcoma-derived cell lines were cultured in a 10 cm dish with full growth medium containing RPMI 1640 supplemented with 10 % FBS. The cell incubator was set at 37 °C supplemented with 5% CO2. Cells were then seeded into 6 well plates. After overnight incubation in the full growth medium, the old medium was replaced with 1.5 ml of RPMI 1640 (supplemented with 10 % dialyzed FBS) containing 11.1 mM [U-^13^C]-glucose or 2 mM [U-^13^C]-glutamine. After 3 hours of ^13^C tracing, intracellular metabolites were harvested.

### Animal Models

All animal procedures were approved by the Institutional Animal Care and Use Committee (IACUC) at Duke University. The mouse model of soft-tissue sarcoma was generated on a mixed background (129/SvJae and C57BL/6) using a combination of alleles that were previously described: Pax7^CreER-T2 20^, p53^FL/FL^ ^21^, LSL-Nras^G12D^ ^22^ and ROSA26^mTmG^ Primary mouse soft tissue sarcomas were generated in the mouse hind limb as previously described ^19^ by intramuscular (IM) injection of (Z)-4-hydroxytamoxifen (4-OHT). 4-OHT was dissolved in 100% DMSO at a concentration of 10 mg/ml and 50 μl of the solution was injected into the gastrocnemius muscle.

### In vivo ^13^C glucose and glutamine infusions

To perform in vivo nutrient infusions, chronic indwelling catheters were placed into the right jugular veins of mice, and animals were allowed to recover for 3-4 days prior to infusions. Mice were infused with [U-^13^C]-glucose for 3 hours at a rate of 20 mg/kg/min (150 µl/hour). Blood was collected via the tail vein at 0, 30 min, 1, 1.5, 2, 2.5, and 3 hours. The plasma was collected by centrifuging blood at 3,000g for 15 min at 4 °C. At the end of infusions, tissues were snap-frozen in liquid nitrogen and stored at -80 °C for further analyses. [U-^13^C]-glutamine (Cambridge Isotope Laboratories) was infused for 3 hours at a rate of 6 mg/kg/min (200 µl/hour).

### HPLC method

The analysis of metabolites in mouse tissues and plasma was performed using Ultimate 3000 UHPLC (Dionex), while the analysis of metabolites from cultured cells was performed using Vanquish UHPLC (Thermo Fisher Scientific). A hydrophilic interaction chromatography method (HILIC) with an Xbridge amide column (100 x 2.1 mm i.d., 3.5 μm; Waters) was used for compound separation at 25 °C. Mobile phase A: water with 5 mM ammonium acetate (pH 6.8), and mobile phase B: 100 % acetonitrile. Linear gradient is: 0 min, 85% B; 1.5 min, 85% B; 5.5 min, 35% B; 6.9 min, 35% B; 10.5 min, 35% B; 10.6 min, 10% B; 12.5 min, 10% B; 13.5 min, 85% B; 17.9 min, 85% B; 18 min, 85% B; 20 min, 85% B. Due to the instrumentation difference between Ultimate 3000 UHPLC and Vanquish UHPLC, different flow rates were used. For Ultimate 3000 UHPLC, the flow rate is: 0-5.5 min, 0.15 ml/min; 6.9-10.5 min, 0.17 ml/min; 10.6-17.9 min, 0.3 ml/min; 18-20 min, 0.15 ml/min. For Vanquish UHPLC, the flow rate is: 0-5.5 min, 0.11 ml/min; 6.9-10.5 min, 0.13 ml/min; 10.6-17.9 min, 0.25 ml/min; 18-20 min, 0.11 ml/min.

### Mass Spectrometry

The analysis of metabolites in mouse tissues and plasma was performed using Q Exactive Plus mass spectrometer (Thermo Fisher Scientific), while the analysis of metabolites from cultured cells was performed using Orbitrap Exploris 480 mass spectrometer (Thermo Fisher Scientific). Both mass spectrometers are equipped with a HESI probe and operated in the positive/negative switching mode. When Q Exactive Plus mass spectrometer was used, the relevant parameters are as listed: heater temperature, 120 °C; sheath gas, 30; auxiliary gas, 10; sweep gas, 3; spray voltage, 3.6 kV for positive mode and 2.5 kV for negative mode; capillary temperature, 320°C; S-lens, 55. The resolution was set at 70,000 (at *m/z* 200). Maximum injection time (max IT) was set at 200 ms and automated gain control (AGC) was set at 3 × 10^6^. When Exploris 480 mass spectrometer was used, the relevant parameters are as listed: vaporizer temperature, 350 °C; ion transfer tube temperature, 300 °C; sheath gas, 35; auxiliary gas, 7; sweep gas, 1; spray voltage, 3.5 kV for positive mode and 2.5 kV for negative mode; RF-lens (%), 30. The resolution was set at 60,000 (at *m/z* 200). Automatic maximum injection time (max IT) and automated gain control (AGC) were used.

### Metabolite extraction from cultured cells, tissues and plasma

To harvest intracellular metabolites, cells were briefly washed with ice cold saline (0.9% NaCl, 1ml, twice) and immediately placed on dry ice before they were extracted into 1 ml extraction solution composed of 80% methanol/water (pre-cooled in -80 °C freezer). Samples were centrifuged at 20,000 g for 10 min at 4 °C, and the supernatant was split into two Eppendorf tubes before drying in a speed vacuum concentrator (Labcono). The dry pellets were reconstituted into 30 μl sample solvent (water: methanol: acetonitrile, 2:1:1, v/v) and 3 μl was injected into the LC-HRMS. The tumor sample was first homogenized in liquid nitrogen and then 5 to 10 mg was weighed in a new Eppendorf tube. Ice cold extraction solvent (250 μl) was added to the tissue sample, and a pellet mixer was used to further break down the tissue chunk and form an even suspension, followed by the addition of 250 μl to rinse the pellet mixer. After incubation on ice for an additional 10 min, the tissue extract was centrifuged with a speed of 20,000 g at 4 °C for 10 min. The dry pellets were reconstituted into 30 μl (per 3 mg tissue) sample solvent (water:methanol: acetonitrile, 2:1:1, v/v), and 3 μl was injected to LC-HRMS. 5 μl mouse plasma was mixed with 5 μl water, and 40 μl ice cold methanol was added. After vortex for 1 min, the mixture was centrifuged with a speed of 20,000 g at 4 °C for 10 min, and 3 μl was injected to LC-HRMS.

### Data analysis and statistics

LC-MS peak extraction and integration were performed using commercially available software Sieve 2.0 (Thermo Fisher Scientific). The integrated peak area was used to calculate ^13^C enrichment. Natural abundance correction was performed using software R with Bioconductor R package IsocorrectionR ^23^. All data are represented as mean ± SD. All p values are obtained from the student’s t-test two-tailed using GraphPad Prism 8 unless otherwise noted.

## Acknowledgments

We thank Clay Rouse, DVM for consultation with the jugular vein surgical procedure. Mice were generously provided by Anton Berns (p53^FL^) and Chen-Ming Fan (Pax7^CreER-T2^). We thank North Carolina State University (NCSU) for financial support. We thank METRIC staff at NCSU for instrument maintenance. This work was supported by the National Institutes of Health R35CA197616 (D.G.K.) T32CA093240 (D.E.C.), Duke Cancer Institute Pilot Grant P30 CA014236 (D.G.K), a grant from the Slifka Foundation and Wendy Walk Foundation (D.G.K).

## Author contributions

D.E.C. performed all animal experiments. X.L. and L.D. performed *in vitro* experiments and LC-MS analysis. X.L., L.D., D.E.C., J.W.L, and D.G.K. participated in experimental design. X.L, L.D., and G.S. interpreted results and wrote the manuscript. All authors provided input on the manuscript.

